# Leopard-EM: An extensible 2DTM package to accelerate in situ structural biology

**DOI:** 10.1101/2025.08.26.672452

**Authors:** Matthew D. Giammar, Joshua L. Dickerson, Laina N. Hall, Bronwyn A. Lucas

**Affiliations:** Center for Computational Biology, University of California Berkeley, Berkeley, CA, USA; Department of Molecular and Cell Biology, University of California Berkeley, Berkeley, CA, USA; California Institute for Quantitative Biosciences (QB3), University of California, Berkeley, CA, USA; Biophysics Graduate Program, University of California, Berkeley, CA, USA; Molecular Biophysics and Integrated Bio-Imaging, Lawrence Berkeley National Lab, Berkeley, CA, USA

## Abstract

The ability to generate high-resolution views of cells with cryogenic electron microscopy (cryo-EM) can reveal the molecular mechanisms of biological processes in their native cellular context. The revolutionary impact of this strategy is limited by the difficulty of accurately annotating structures within these images. 2D template matching (2DTM), in which high-resolution structural models are used as computational probes to locate and orient molecular complexes with high precision, has shown initial promise in annotating single molecules in cellular cryo-EM images. While the scientific community works to identify best practices for applying 2DTM to specific biological questions and to maximize sensitivity and throughput, a modular and extensible software architecture would support the rapid development of novel methodological approaches, thus accelerating innovation within the field. To achieve this, we developed Leopard-EM (**L**ocation & ori**E**ntati**O**n of **PAR**ticles found using two-**D**imensional t**E**mplate **M**atching), a modular Python-based 2DTM implementation built to be easily customizable. We implemented an automated pixel size refinement procedure and find that 2DTM is sensitive to pixel size to within 0.001Å. To demonstrate the flexibility of the Leopard-EM architecture, we developed a constrained search protocol that improved small ribosomal subunit (SSU) detection by approximately eightfold by using initial locations and orientations determined for the large ribosomal subunit (LSU). Using this strategy, we captured a distribution of ribosome rotation states within a living cell at single-molecule resolution. We envision that Leopard-EM can be used as a platform for development of *in situ* cryo-EM data processing workflows, facilitating the rapid development of this field. Leopard-EM is available at https://github.com/Lucaslab-Berkeley/Leopard-EM.

## 1 Introduction

Cryogenic electron microscopy (cryo-EM) can generate micrographs of cells, each containing a large number of biological macromolecules in their native cellular environment. Reliable annotation of specific biological macromolecules within these images is the rate limiting step of *in situ* structural biology. In contrast to single-particle analysis (SPA), in which particle picking methods use high-contrast, low-resolution features, many cellular complexes are visually indistinguishable relative to the cellular milieu. Moreover, cell sections are typically 100-200 nm in thickness, resulting in a higher proportion of multiple and inelastic scattering, reducing signal-to-noise ratio (SNR) relative to SPA. These limitations, combined with the inherently low SNR of cryo-EM images due to radiation damage and the low-dose collection scheme, make the specific annotation of even a handful of the ∼20,000 proteins in the proteome an extreme challenge.

Improved accuracy in structure prediction has made it possible to generate a reference ‘structure-ome’ from genomic sequences [14, 2, 34, 16]. The greatly increased number of available structural models has made template matching a viable strategy to locate proteins in cryo-EM images of cells. Template matching in cryo-EM refers to several strategies in which a reference is used to detect a particular feature of interest by cross-correlation with the data. Template matching strategies vary in the type of template used, the inclusion and type of noise applied to the template, the number of alternative templates considered, the filters applied, and the way in which the results are normalized. Each of these differences changes the interpretation of the results and makes a quantitative comparison challenging.

Even in crowded cellular environments, high-resolution structural features remain distinct and are best preserved in full-exposure images of untilted samples, rather than in tomograms [8]. 2D dimensional template matching (2DTM), by exploiting these high-resolution features, can be used to determine the location and orientation of macromolecules in cryo-EM images with high precision, even in the presence of background noise [31, 28]. In 2DTM, the presence of a particular complex is assessed by calculating the cross-correlation of spectrally whitened projections, corresponding to specific poses of a noise-free 3D reference template, with the whitened image. The cross-correlation of the best matching orientation at each pixel is stored, as well as the rotational offset to generate the projection from the template. Matches are detected by thresholding based on the probability of a false detection given Gaussian noise and the total number of cross-correlations performed. 2DTM is, in essence, an implementation of the classical matched filter [31], which is the theoretically optimal linear filter for the detection of a signal of interest in the presence of additive random noise.

2DTM has been applied to identify the location and orientation of VP1 polymerase in rotavirus DLP particles [28], ribosomes in thin extensions of mammalian cells [26, 27], as well as ribosomes and ribosomal subunit precursors in *Mycoplasma pneumoniae* [19] and FIB-milled yeast lamellae [20].Similar strategies have been applied to locate particles of interest for subsequent high-resolution 3D reconstruction from 2D images [4, 18], and have generated high-resolution *in situ* structures of translating ribosomes [5, 38] and large photosynthetic supercomplexes [35].

2DTM has the potential to visualize the proteome in context, but further improvements in sensitivity and throughput are required to achieve this goal. In addition, customization is required to address specific biological questions or datasets. For example, comparing 2DTM z-scores from multiple templates was used to discriminate structurally related large ribosomal subunit (LSU) maturation intermediates within the yeast nucleus [20]. In another example, mammalian translation dynamics were estimated by ordering ribosome-cofactor complexes on the basis of their conformational state [27]. Alternate metrics to call significant detections have also been proposed [36]. Several extensible software packages have been developed for 3D template matching (3DTM) [22, 21, 3], providing frameworks for flexible methodological development in that domain. However, comparable modular and customizable implementations for 2DTM are lacking.

Realizing this potential requires a modular and accessible software architecture that can be easily adapted by various users to fit the application at hand. Here we describe a new modular implementation of 2DTM, Leopard-EM, which is written in Python, leveraging PyTorch for GPU acceleration. This tool was designed to be extensible for rapid development by a broader user and developer base. We show how Leopard-EM can be used in a standard 2DTM workflow and demonstrate the importance of screening template pixel size to maximize detection sensitivity. We also demonstrate the extensibility of Leopard-EM by presenting a use case in which results from a prior template matching search for the large ribosomal subunit are used to constrain a local search for the small ribosomal subunit, enabling identification of intersubunit rotation with single-molecule precision.

## 2 Results

### 2.1 Leopard-EM is an extensible, GPU-accelerated Python package for 2DTM

To enable the adaptation of 2DTM to a wider range of biological problems, we have implemented the 2D template matching algorithm [28, 19, 27] in a flexible and extensible Python package. Python has become the *lingua franca* for scientific programming and machine learning thanks to a rich infrastructure for data science and performant numerical libraries [12, 23, 24]. Developing this 2DTM package within the Python ecosystem benefits from this preexisting infrastructure and enables easy integration into larger cryo-EM workflows through Python scripts. We designed this 2DTM package with modularity in mind, separating user-facing programs, data validation and preprocessing steps, and core backend functionality into three different levels (Figure 1). We named this Python package Leopard-EM, for **L**ocation & ori**E**ntati**O**n of **PAR**ticles found using two-**D**imensional t**E**mplate **M**atching.

**Figure 1.**
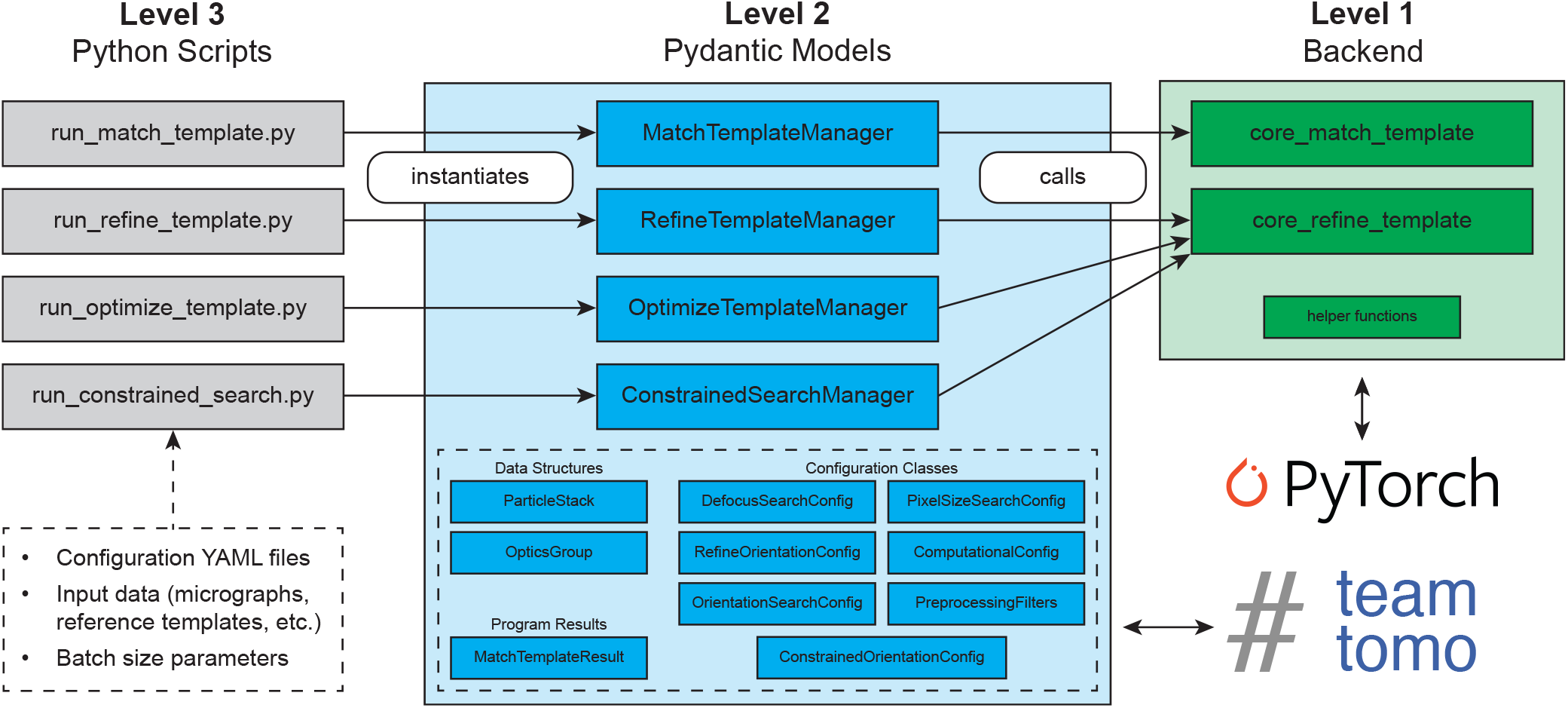
Overview of the multi-level Leopard-EM package. The third level of Leopard-EM are pre-written Python scripts (gray) for a particular program which instantiate a manager object at the second level (blue). Each manager object uses other Pydantic models, which encapsulate common data structures, define validation methods for inputs, and act as connectors to cryo-EM data processing methods defined in TeamTomo, to configure program inputs in a hierarchical manner. Manager objects call one of the backend functions in the first level (green) to run a program. The backend of the Leopard-EM package leverages PyTorch to interface with computer hardware, namely GPUs, to accelerate the computationally intensive stages of 2DTM.

The core of the 2DTM method is the cross-correlation of two-dimensional projections from a reference model with a high-resolution cryo-EM micrograph [19]. Calculating these cross-correlations greatly benefits from GPU hardware, and we implemented this core 2DTM functionality using PyTorch primitives for accelerated GPU computation. The computational backend of Leopard-EM is designed with a simple, data-oriented interface which expects an explicit definition of the search space, including a set of 3D orientations (e.g., Euler angles) and a set of contrast transfer function (CTF) parameters used to search across different defocus planes. Each point in this search space corresponds to a specific combination of orientation and defocus which are used to generate two-dimensional projections from a 3D reference template. By keeping the backend simple and separating it from the input parsing or preprocessing steps, we allow the possibility of using more performant backends other than PyTorch in the future without major modifications to the user-facing portions of the Leopard-EM package.

One layer up from the backend module, we included a set of custom classes built on Pydantic [6], a data validation library for Python, to facilitate input configuration, data parsing, and results export in Leopard-EM (Figure 1). These Python classes model input and output data structures, implement preprocessing steps, define the 2DTM search space, and dispatch the computation to the backend module. By adopting this object-oriented design, we reduce overall complexity by reusing common structures and methods as well as moving complicated, single-time preprocessing steps out of the backend. This modular framework will enable rapid development of 2DTM workflows.

The final layer of Leopard-EM is the user-facing interface which consists of easily configurable Python scripts for running Leopard-EM programs. Each program is configured using a YAML file, and the contents of the YAML file are mapped to Leopard-EM Pydantic models. Example YAML configurations and Python scripts can be found on the Leopard-EM GitHub page, and the configuration of each program is discussed in the online Documentation (lucaslab-berkeley.github.io/Leopard-EM/). Working closely with TeamTomo (teamtomo.org), a collaborative open source project for cryo-EM infrastructure in Python, during the development of Leopard-EM has allowed us to focus on the 2DTM algorithm and reduce the time spent re-implementing common cryo-EM data processing primitives. Specifically, we used the Fourier slice projection operation from `torch-fourier-slice` in the core 2DTM algorithm, and contributed packages to simulate electron scattering maps (`ttsim3D`), sampling points in orientation space (`torch-so3`), and generating Fourier filters including the CTF (`torch-fourier-filter`). By making these primitive operations available as standalone Python packages, we hope to simplify future developments of Python-based cryo-EM workflows.

Adopting this modular framework makes Leopard-EM easy to extend, enabling future enhancements to both the speed and sensitivity of 2DTM as well as custom workflows to address specific biological questions.

### 2.2 Leopard-EM can locate and orient different macromolecules through exhaustive searches

In the standard 2DTM workflow, we first run the program `match_template`, which performs exhaustive orientation and defocus sampling to locate and orient particles within a micrograph. This program generates and cross-correlates reference template projections with the micrograph across all specified orientations and sample defocus planes. The best matching orientation and defocus, based on the highest cross-correlation value, is iteratively updated on a per-pixel basis as the program goes through the search space. After searching for all orientations and defoci, the maximum cross-correlation scores at each pixel are compared against a Gaussian noise model, and positive detections are assigned using a threshold chosen based on the expected number of false positives per micrograph (which is 1 by default)[28]. The orientation and defocus assignments are then further refined using the program `refine template` which performs a local search around previously identified particles by sampling a finer grid of orientations and defocus values. The more precise results of `refine template` can then be used for further downstream processing, such as visualizing the locations and orientations of identified molecules or 3D reconstruction.

We confirmed that we can use the standard 2DTM workflow in Leopard-EM to detect single ribosomes in cryo-EM images of yeast lamellae, as previously demonstrated [19, 27, 20]. We applied Leopard-EM to independently detect large (LSU) and small (SSU) ribosomal subunits in cryo-EM images of yeast cell cytoplasm (Figure 2A,B,C). Notably, the detected LSU and SSU particles align in space (Figure 2D,E), and their orientations are consistent with the formation of 80S ribosomal complexes (Figure 2F,G,H), providing further validation of the significance of these detections.

**Figure 2.**
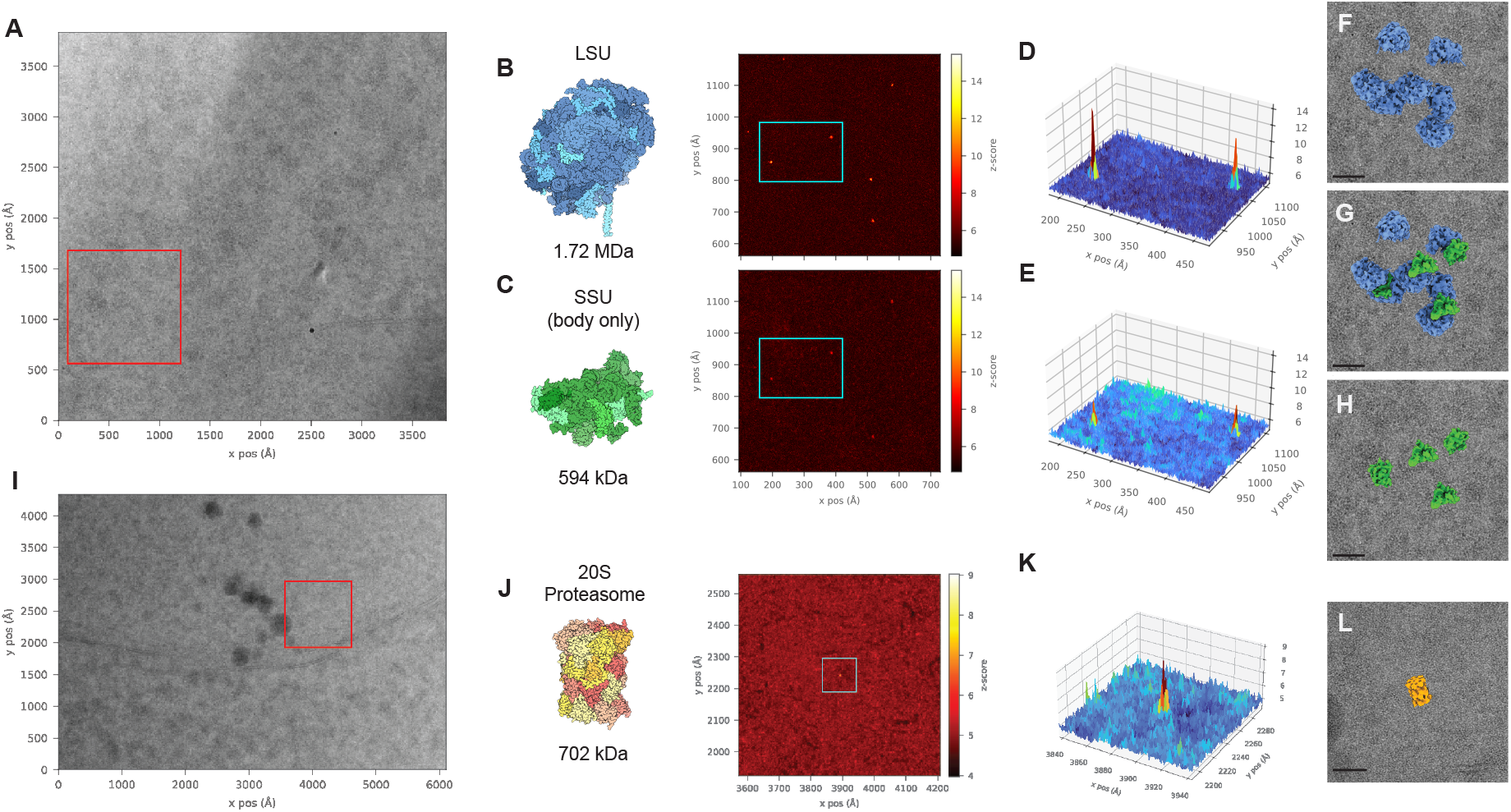
Representative two-dimensional template matching (2DTM) results for searches run using a large ribosomal subunit (LSU), small ribosomal subunit (SSU) body, and 20S proteasome as reference templates. (**A**) Micrograph used for both LSU and SSU body 2DTM searches collected at 0.936 Å/px. (**B**) (left) LSU template with simulated molecular mass below. (right) 2DTM z-scores for LSU in the region of interest (red box in **A**). (**C**) (left) SSU template with simulated molecular mass below. (right) 2DTM z-scores for SSU in the region of interest (red box in **A**). (**D,E**) Zoomed-in z-score surface maps within cyan box for the LSU and SSU, respectively. (**F,G,H**) The relative position and orientations of LSU and SSU templates overlaid on the micrograph region of interest where **F** shows the LSU alone, **G** shows both the LSU and SSU, and **H** shows the SSU alone. (**I**) Micrograph used for the 20S proteasome 2DTM search collected at 1.06 Å/px. (**J**) (left) Proteasome template with simulated molecular mass below. (right) 2DTM z-scores for the proteasome search in the region of interest (red box in **I**). (**K**) Z-score surface map for the identified proteasome peak. (**L**) 20S proteasome template at identified location and orientation overlaid on the region of interest. Scale bars in **F, G, H**, and **L** correspond to 20 nm.

Ribosomes are large, abundant, and primarily consist of RNA, making them relatively prominent features in cellular cryo-EM images and presumably easier to detect with 2DTM. We show that we can also apply Leopard-EM to detect the protein-only 20S proteasome (Figure 2I-L). We found 14 significant detections in 7 of the 28 images searched. Although we expect many detections to be missed, we only found significant detections of the proteasome in the nucleus close to the nuclear periphery (Figure 2I), consistent with a previous report of enrichment at this cellular location in *Chlamyodomonas reinhardtii* [1].

#### 2.2.1 Pixel size refinement is essential for optimal detection of complexes with 2DTM

2DTM provides a read-out of the agreement between a structural model and the image. For accurate detection, it is essential that the pixel sizes of both the model and the images are correct. Any discrepancy causes the template’s size and interatomic distances to differ from the target molecule (Figure 3), reducing the cross-correlation score and lowering the 2DTM sensitivity. However, the extent of this effect has not been systematically evaluated.

**Figure 3.**
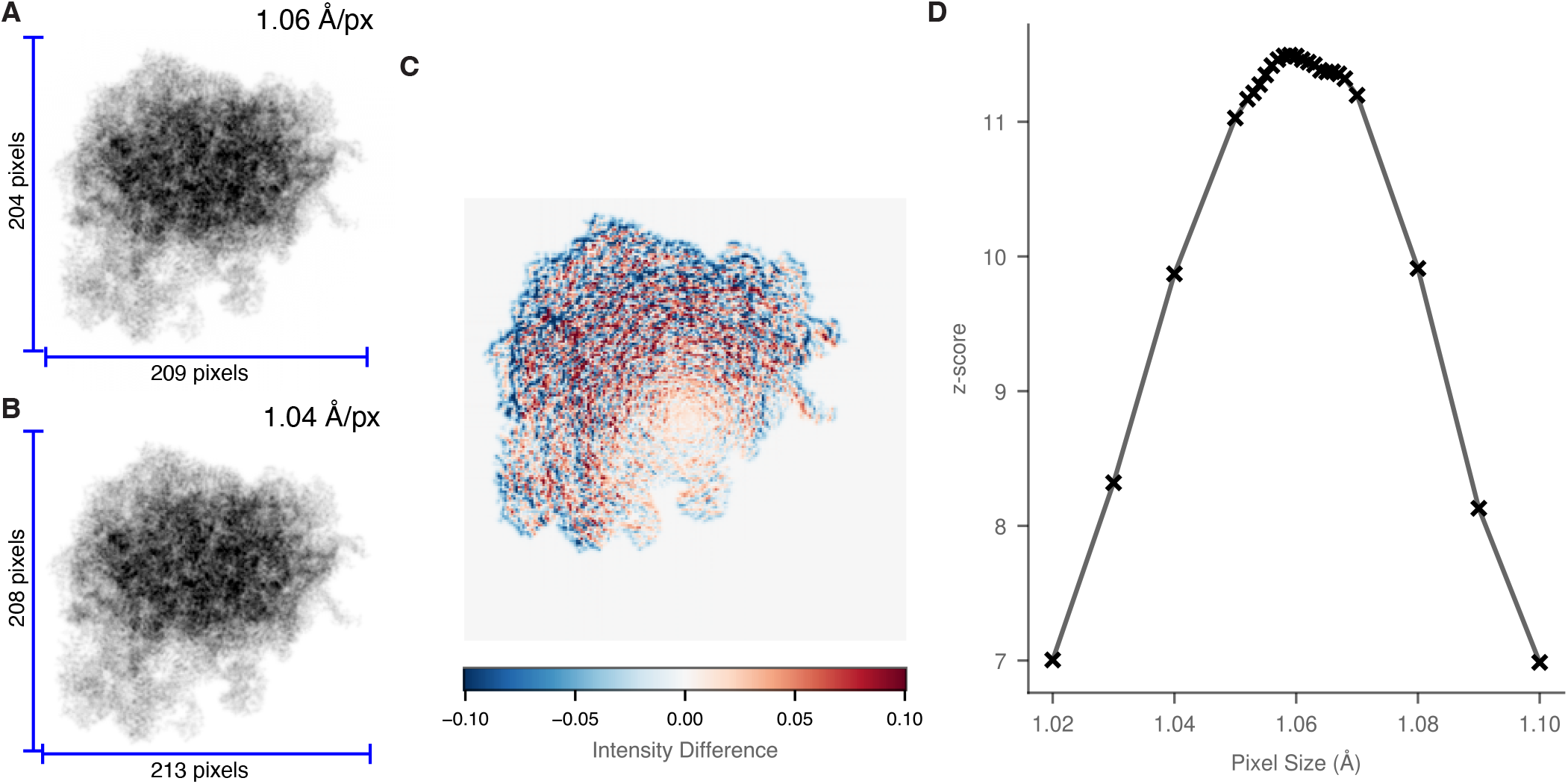
2DTM is sensitive to pixel size errors of < 1%. (**A,B**) Projection images from the same orientation of the LSU template with different with pixel sizes, 1.06 Å (**A**) or 1.04 Å (**B**). A < **2**% difference causes significant changes in the size of template projections. (**C**) Intensity plot showing the difference in the intensity between (**A**) & (**B**), showing a large change in intensity from 1.06 Å to 1.04 Å. This causes significant differences in the 2DTM z-scores. (**D**) Plot showing the change in z-score of a set of six LSU detections with changing template pixel size. The difference in pixel size shown in (**A,B**) causes a 15% decrease in 2DTM z-scores.

To address this, we implemented a template pixel size optimization step in Leopard-EM through the `optimize template` program. This requires a pre-identified set of particles, which can be quickly obtained by running template matching on a cropped image. The optimization uses the package `ttsim3d` to simulate 3D volumes of the model at different pixel sizes, then cross-correlates projections of these from the annotated orientations with the image. The optimal pixel size is determined by maximizing the mean z-score of the top *N* peaks, where *N* is a user-definable parameter. We note that this method simply scales the pixel size of the template to match the micrograph and is therefore agnostic to whether errors are with the model or micrograph pixel size. Therefore, it can only be used for microscope magnification calibration if the model pixel size is reliable, which is often true for X-ray-derived models but less certain for those built into cryo-EM maps.

We used images from a previously published dataset [17] to test the effect of pixel size on 2DTM z-score. Our results show that 2DTM is highly sensitive to pixel size, with detectable effects at differences as small as 0.001 Å (Figure 3D). The change in z-score was not symmetric, with a sharper penalty when the pixel size was underestimated relative to when the pixel size was overestimated. At this stage, it is unclear whether this is a function of specific features in the LSU template or a more general feature of the method. An error of just 0.02 Å (less than 2%) results in a 15% drop in z-score. Since SNR scales with the square root of molecular mass [13, 28], this drop in SNR corresponds to a ≈40% increase in the minimum detectable molecular mass, making it significantly harder to identify small macromolecules. Considering that pixel size errors exceeding 1% are frequent in cryo-EM datasets [7], the pixel sizes of the model and image cannot be assumed to align. We therefore recommend performing pixel size calibration for each combination of imaging conditions and template model, which can be done using the `optimize template` program in Leopard-EM.

### 2.3 A constrained search as an example of extensibility

To demonstrate the extensibility of Leopard-EM, we incorporated a constrained search that uses the locations and orientations of one type of macromolecule to constrain the search for another. The identification of macromolecules *in situ* using cryo-EM is fundamentally limited by the low-dose conditions required due to radiation damage. As a result, 2DTM is currently limited to detecting proteins with a molecular mass of at least 300 kDa *in situ* under ideal conditions [27]. In 2DTM, the incorporation of high spatial frequency information creates narrow cross-correlation peaks with respect to orientation and defocus, which in turn requires fine angular and defocus sampling to reliably capture those peaks. However, this fine sampling results in a high noise floor caused by a large number of cross-correlations, necessitating a high threshold. With more independent comparisons, the chance of obtaining a high z-score purely by random chance also increases, thus raising the effective noise floor and requiring a more stringent detection threshold. Reducing the number of cross-correlations directly reduces the probability of observing high z-scores due to noise alone, and therefore permits a lower detection threshold for the same false positive rate [28]. This can be achieved by restricting the search space in orientation (*ϕ, θ, ψ*) or position (*x, y, z/*defocus) (Figure 4A). For example, if both the location and orientation of a particle are known in advance, the reduced number of comparisons can lower the detection threshold sufficiently to detect particles an order of magnitude smaller at a false-positive rate of 1 in 200 particles.

**Figure 4.**
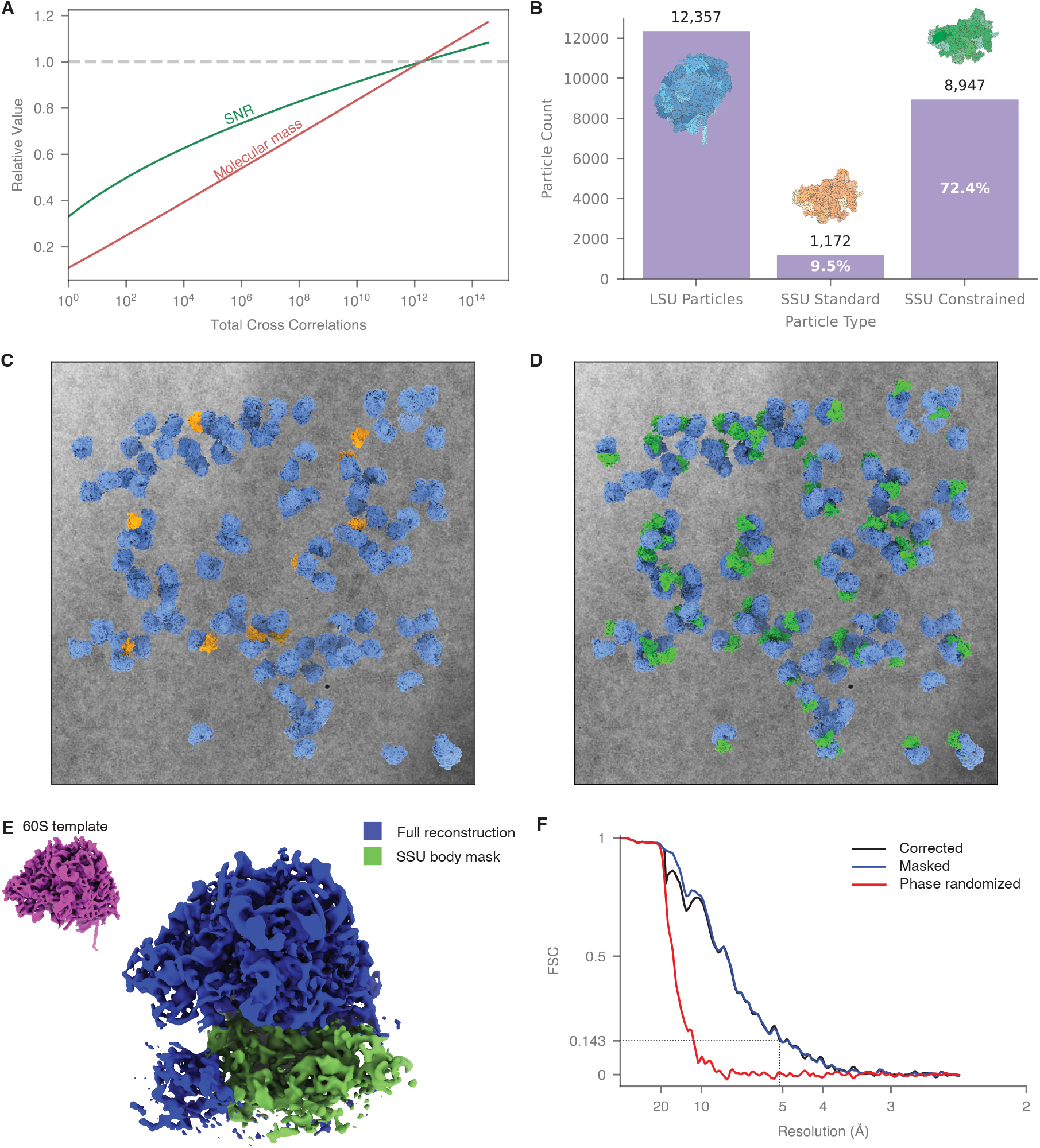
Constrained searches improve the sensitivity and functional resolution of 2DTM. Constraining the search space in 2DTM — by limiting the number of cross-correlograms (CCG) through known molecular positions and orientations — lowers the signal-to-noise ratio (SNR) threshold for detection, thereby enabling identification of smaller molecular complexes. (**A**) Plot showing the minimum detectable molecular mass at a false-positive rate of 1 in 200 particles, relative to the standard threshold of 1 false positive per micrograph. When both position and orientation are known, the detection threshold improves by approximately 10-fold. (**B**) Plot showing the number of detections in 62 micrographs when performing an unconstrained search for the LSU and SSU body or constraining the search for the SSU body using positions and orientations from the LSU search (right). (**C**) Simulated slab of LSU particles (blue) and SSU particles (orange) found using the unconstrained 2DTM search overlaid on a representative micrograph. (**D**) The same LSU particles and micrograph now depicted with the constrained search SSU particles (green) showing that the constrained search increases 2DTM sensitivity. (**E**) Reconstructed volume of the LSU particles where a SSU particle had also been detected. The SSU is clearly present in the reconstruction despite not being present in the template. To estimate the resolution, a mask was created around the SSU body (green). This region reconstructed to 5.1 Å, as determined by a Fourier Shell Correlation of 0.143(**F**)[29].

Beyond enabling the detection of smaller proteins, constrained searches also reduce false negatives when applied to larger molecules. We implemented this strategy in Leopard-EM by allowing users to restrict the search space via symmetry specifications or explicit angular ranges within the `match_template` program. Additionally, we developed a separate program to perform constrained searches on particle stacks, using known positions and orientations of one molecule to guide the detection of another. This program, `constrained search`, can serve as a model for others looking for customization within the 2DTM framework.

#### 2.3.1 Constrained searches identify small ribosomal subunits

We demonstrate this approach by searching for the ribosome SSU body, which has a molecular mass in the template of approximately 594 kDa. When performing a full unconstrained search, we detected 1,172 SSU particles in a dataset of 62 micrographs, compared to 12,357 LSU particles. Using the positions and orientations derived from the LSU 2DTM results, the constrained search yields significant SSU detections adjacent to 72% of LSUs, an ≈ 8-fold increase (Figure 4B). An example micrograph overlaid with these results reprojected is shown in Figure 4C–D. We used the 2DTM locations and orientations to reconstruct LSUs where we had detected an associated SSU and recovered the density of the SSU, although it was not present in the template (Figure 4E). The SSU reconstruction reached an estimated resolution of 5.1 Å (Figure 4F), determined using a mask encompassing only the SSU body, which excluded the original LSU template to avoid template bias [18].

#### 2.3.2 2DTM can be used to characterize a continuous distribution of rotated small ribosomal subunits

The constrained search also enables us to capture continuous conformational heterogeneity at singlemolecule resolution. The ribosome is a dynamic molecular machine comprising two subunits, an LSU and an SSU, which rotate relative to each other during mRNA translation and peptide synthesis [15]. By comparing structural models determined in different rotational states, we constrain the search to one or two rotation axes, corresponding to intersubunit rotation (Ψ offset) and roll (*θ* offset) [32]. We find a predominate population with less than 2.5° Ψ offset relative to the non-rotated ribosome template [33]. The remaining approximately 25% of the ribosomes are in a rotated conformation (Figure 5A), consistent with previous findings [27]. We also find that there is a small rotation on the orthogonal axis, consistent with SSU body roll (Figure 5B), which coincides with the rotated state (Figure 5C). To validate these results, we generated 3D reconstructions of LSUs classified as associated with SSUs that are at least 4° rotated or less than 4° rotated, which could clearly be distinguished (Figure 5D).

**Figure 5.**
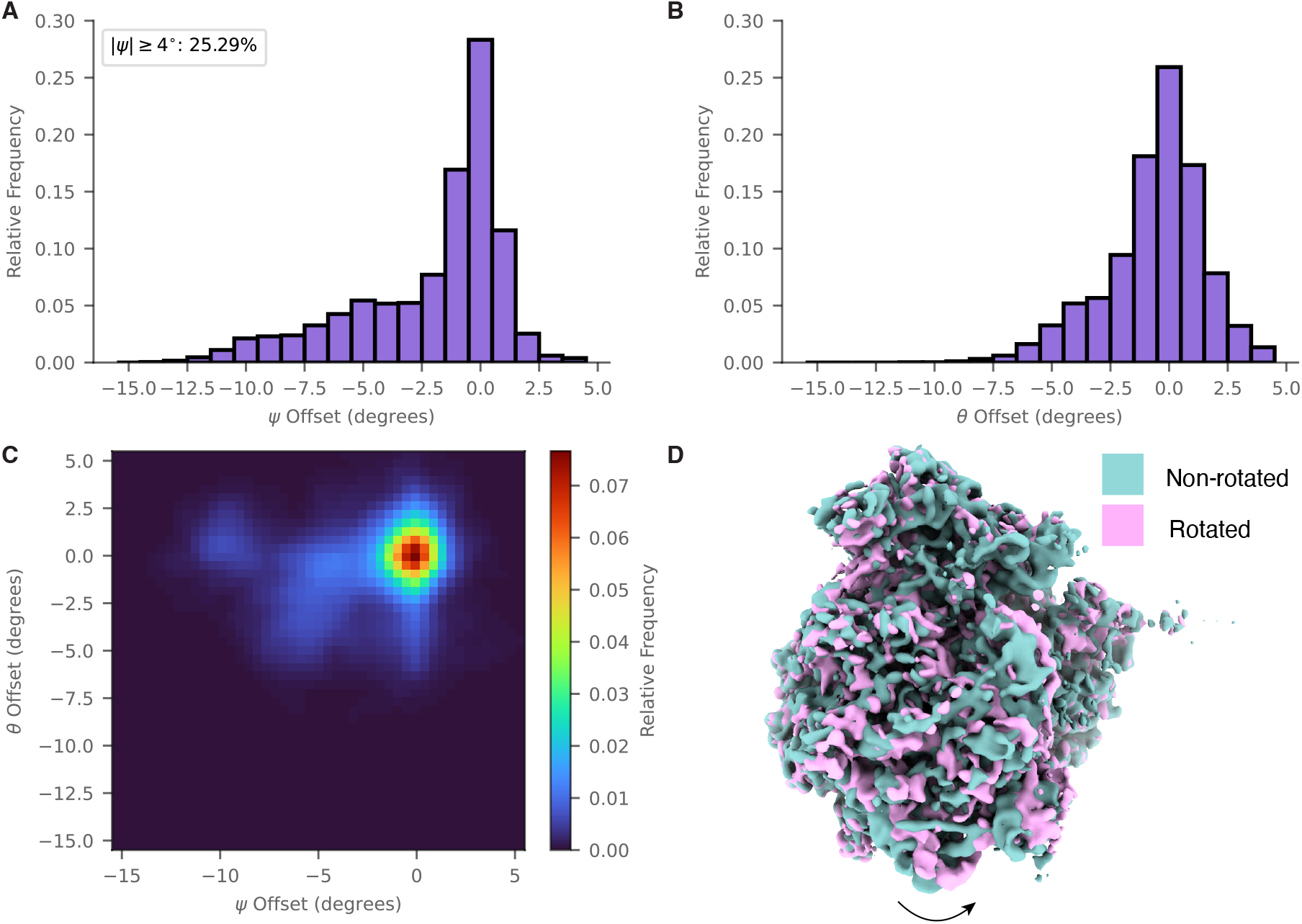
Constrained searches allow identification of a continuous distribution of conformational states. (**A**) A histogram showing the angular distribution of SSU rotations relative to the LSU, as determined by constrained 2DTM. (**B**) A histogram showing the angular distribution of SSU at a secondary axis (subunit roll) relative to the LSU, as determined by constrained 2DTM. (**C**) A 2D heatmap combining the results shown in (**A**) and (**B**) where the intensity corresponds to the occupancy of SSUs. (**D**) Reconstructions of LSU particles, classified by the output of 2DTM as having a rotated (≥ 4°) or non-rotated (< 4°) SSU.

## 3 Conclusions and Future Perspectives

We have developed a new software package for 2DTM, Leopard-EM, which has been written with extensibility at the forefront. By implementing the package in Python and in a modular framework, we make this approach more accessible to the cryo-EM community and allow new features to be implemented more rapidly. This is crucial in a fast-moving field, where small improvements in sensitivity and speed can open up new areas of investigation. We demonstrate that the standard 2DTM workflow can be used in Leopard-EM to identify and align RNA containing ribosomes and macromolecules without RNA. The main advantage of Leopard-EM lies in the ability to quickly implement non-standard data processing strategies, which we demonstrate by incorporating a constrained search and using it to increase sensitivity and identify continuous heterogeneity in SSU rotation states.

The sensitivity of 2DTM is currently limited to the detection of large complexes greater than 300 kDa in the best-case scenario. This limitation is largely due to the difficulty in discriminating true from false positives. By constraining the search for the small ribosomal subunit using prior information from the large ribosomal subunits, we reduced the noise level, enabling us to distinguish true positives from false positives at lower SNR. Using this strategy, we recovered ≈ 8 × more SSU detections, demonstrating the potential of using prior information to improve the detection of smaller particles. This approach could easily be extended to incorporate other types of prior information, such as membrane features or other visible cellular structures.

We show that 2DTM can characterize molecular motions. During translation, the orientation of the SSU relative to the LSU changes. By performing a constrained local search for the SSU using two rotation axes defined on the LSU, we defined the distribution of rotation and roll states with single-molecule precision, without the need for 3D reconstruction and classification, consistent with reference [27].

Leopard-EM was designed to make it easy to develop custom workflows and new functionality. Structural cell biology is an emerging field and the role of untilted data collection and 2DTM are still being explored. Possible future improvements include substituting PyTorch for a custom, faster back-end, application of custom Fourier filters, development of new scoring metrics, and combining untilted with tilted data collection. Other modifications will likely emerge as this approach finds new applications.

The development of cryo-EM has always been dependent on software tools. Leopard-EM benefits from the modular tools that have been developed as part of TeamTomo that can be adapted to fit the question at hand. We expect that reducing the barrier to entry will enable a broader developer base to contribute to the development of new strategies to further improve single-molecule structural biology in cells.

## 4 Code and data availability

The code described in this manuscript can be found at the Leopard-EM GitHub page for version v1.0. The continuous development of the package can be followed at the Leopard-EM main GitHub page.

Data used throughout this manuscript will be published in a database before publication. Scripts for processing the data and generating the figures can be found at the Leopard-EM manuscript GitHub page.

## 5 Methods

### 5.1 Yeast cell culture, grid preparation and FIB-milling

*Saccharomyces cerevisiae* strain W303 haploid (NucLoc) colonies were inoculated in 5 mL of YPD media and grown overnight at 30°C with shaking (200 RPM). Cultures in mid-log phase (≈ 0.8 OD) were selected and diluted to 10,000 cells/mL. 3 µL were applied to a Quantifoil 2/2 SiO2 200 mesh Cu grid, allowed to rest for 15 s, back side blotted for 8 s at 27°C, 95% humidity, and plunge-frozen in liquid ethane at –184°C using a EM GP2 cryo-plunger (Leica). Frozen grids were stored in liquid nitrogen until FIB-milling.

### 5.2 FIB Milling

Lamella were generated using a Hydra cryo-FIB-SEM (Thermo Fisher) and a Xenon ion source operated at 30 kV. Lamella were processed using AutoTEM(Thermofisher) using the following protocols: Stress relief cuts were milled at a current of 0.3 nA. Rough milling was performed at 0.3 nA. Medium milling was performed at 0.1 nA using a cleaning cross-section pattern, followed by finer milling at 60 pA, ultimately resulting in a targeted thickness of ≈ 500 nm. Polishing was performed at 6 pA to a target thickness of 200 nm. Electron images were not taken during lamella preparation.

### 5.3 Cryo-EM data collection and preprocessing

Cryo-EM data were collected following the protocol described in [20] using a Thermo Fisher Krios 300 kV electron microscope equipped with a Falcon 4 camera and Selectris × energy filter at a nominal magnification of 81,000× (pixel size of 0.936 Å^2^) and a 100 µm objective aperture. Movies were collected to a total fluence of 50 e^*−*^/Å^2^ with 1800 frames in EER mode. Cytoplasm was targeted by selecting regions of interest in a low-magnification overview. The movie frames were aligned using MotionCor3 [37] and dose-weighted according to reference [10]. CTF estimation was perfromed using CTFFIND5 [9].

### 5.4 2DTM search for LSU and SSU body

In summary, we used a PDB model of the 80S ribosome in the non-rotated state, 6Q8Y [33], and manually generated two additional PDB files, one from the LSU alone and one from the SSU. The SSU was further processed by removal of GBLP, RS29A, RS28A, RS25A, RS20, RS12, RS31, RS3, RS10A, RS15, RS18A, RS19A, RPS5P, RS16A, RSSA1, RS17A, S19A and nucleotides 1128-1609 of the modeled 18S rRNA to generate the SSU body alone. These models were aligned with the full model in ChimeraX [25]. The 80S model was re-centered at coordinates (0, 0, 0), and the same transformation was applied to the other models using a custom Python script. After optimizing the pixel size to a value of 0.936 Å (§5.8), we simulated a 512 × 512 × 512 pixel model using the program `ttsim3d`, with the parameters of no re-centering, no additional *B* -factor, and *B* -scaling of 0.5 relative to those deposited in the PDB. Only non-hydrogen atoms were included in the simulation. We ran match template in Leopard-EM with uniform angular sampling, with a psi step of 1.5^°^ and theta step 2.5^°^. The CTF *B* -factor was 0 Å^2^. The defocus search range was -1200–1200 Å in 200 Å steps.

After running `match_template`, the peaks were extracted from the z-score map using a threshold of one false-positive per micrograph. The resulting peaks were further refined, using a defocus search of -100–100 Å in 20 Å steps, an extracted box size of 518 × 518 pixels, and angular steps of 0.05^°^. In total, we located 12,357 LSUs from 62 micrographs (Figure 4B).

All scripts used to perform 2DTM can be found at https://github.com/Lucaslab-Berkeley/Leopard-EM_manuscript.

### 5.5 2DTM search for proteasome

We used the PDB model of the 20S yeast proteasome, 1RYP [11], to simulate a 384 × 384 × 384 pixel model at 1.059 Å using the program `ttsim3d` with no *B* -factor scaling or additional *B* -factor. Only non-hydrogen atoms were included in the simulation. We ran `match_template` in Leopard-EM on 28 previously published micrographs [19] with uniform angular sampling using a psi step of 1.5^°^, theta step of 2.5^°^, and a C2 symmetry.

All scripts used to perform 2DTM can be found at https://github.com/Lucaslab-Berkeley/Leopard-EM_manuscript

### 5.6 Constrained search for SSU

To search for the SSU, we simulated a 3D map of the SSU body using the same parameters as for the LSU. We ran `match_template` in Leopard-EM using the same parameters and used the variance from this result for the z-score calculation in subsequent constrained searches. We determined the rotation axis by calculating the rotation needed to transpose the PDB model 3J77 (rotated) onto 3J78 (non-rotated) using the script ‘get rot axis.py’ provided in Leopard-EM. The vector between the center of the LSU and the SSU, which is necessary to adjust the defocus for each particle, was calculated using the script ‘get_center_vector.py’.

The constrained searches were then performed in four steps using the `constrained_search` program in Leopard-EM. By performing sequentially finer searches, we limit the number of cross-correlations, thus lowering the noise floor. First, a search was performed over the Z-axis (*ψ*) in the range -13–2.5^°^ in 1^°^ steps. The second search, using the orientations from the first, was over the Y-axis (*θ*) from -6.0–4.0^°^ in 1^°^ steps. The third search was over both angles, between -5–5^°^ in 0.5^°^ steps. Finally, a search was performed over both angles between -0.5–0.5^°^ in 0.1 ^°^ steps. This final step included a defocus search in the range -100–100 Å in 20 Å steps. The results from the constrained searches were accumulated using the script ‘sequential threshold processing.py’, using a final threshold of one false positive for every 200 particles.

All scripts for the constrained search can be found at https://github.com/Lucaslab-Berkeley/Leopard-EM_manuscript.

### 5.7 3D reconstructions

The Leopard-EM results files were converted to RELION compatible star files for LSU particles with a detected SSU. 3D volumes were generated for all particles, non-rotated particles, and rotated particles, using RELION’s reconstruct program [30]. After low-pass filtering the maps to 30 Å, masks for the full 80S ribosome were generated in RELION. These masks were manually modified in ChimeraX [25] to create a mask for the SSU body. Postproccesing was performed in RELION to generate the masked 3D maps and estimate the resolution. The maps were low-pass filtered to 8 Å for visualization purposes.

All scripts used to generate the 3D reconstructions can be found at https://github.com/Lucaslab-Berkeley/Leopard-EM_manuscript.

### 5.8 Automated pixel-size search

A single micrograph for each dataset was cropped to a size of 1024 × 1024 pixels for use in pixel size calibration. After running `match_template` on the cropped micrograph, the pixel size optimization was performed using the Leopard-EM program `optimize_template`. The simulation parameters were the same as those described in §5.4. The pixel size range was -0.04–0.04 Å in 0.01 Å steps for the coarse search and -0.008–0.008 Å in 0.001 Å steps for the fine search.

The scripts for this can be found at https://github.com/Lucaslab-Berkeley/Leopard-EM_manuscript.

## 6 Acknowledgements

This work was supported by an NIH DP2 to BAL. JLD is supported by a Helen Hay Whitney Postdoctoral Fellowship, BAL is a Searle Scholar and Shurl and Kay Curci Scholar and LNH is supported by the NSF GRFP. We thank Alister Burt for helpful discussions and critical reading of the manuscript.

## 7 Author contributions

MDG: Conceptualization, Methodology, Software, Formal analysis, Investigation, Resources, Data Curation, Writing - Original Draft, Writing - Review & Editing, Project administration

JLD: Conceptualization, Methodology, Software, Formal analysis, Investigation, Resources, Data Curation, Writing - Original Draft, Writing - Review & Editing, Project administration

LNH: Methodology, Resources, Data Curation, Writing - Review & Editing

BAL: Conceptualization, Writing - Original Draft, Writing - Review & Editing, Supervision, Project administration, Funding acquisition, Methodology, Resources, Data Curation, Investigation

## References

[1] S. Albert, M. Schaffer, F. Beck, S. Mosalaganti, S. Asano, H. F. Thomas, J. M. Plitzko, M. Beck, W. Baumeister, and B. D. Engel. Proteasomes tether to two distinct sites at the nuclear pore complex. Proceedings of the National Academy of Sciences, 114(52):13726–13731, Dec. 2017. Publisher: Proceedings of the National Academy of Sciences.

[2] M. Baek, F. DiMaio, I. Anishchenko, J. Dauparas, S. Ovchinnikov, G. R. Lee, J. Wang, Q. Cong, L. N. Kinch, R. D. Schaeffer, C. Millán, H. Park, C. Adams, C. R. Glassman, A. DeGiovanni, J. H. Pereira, A. V. Rodrigues, A. A. V. Dijk, A. C. Ebrecht, D. J. Opperman, T. Sagmeister, C. Buhlheller, T. Pavkov-Keller, M. K. Rathinaswamy, U. Dalwadi, C. K. Yip, J. E. Burke, K. C. Garcia, N. V. Grishin, P. D. Adams, R. J. Read, and D. Baker. Accurate prediction of protein structures and interactions using a three-track neural network. Science, 373(6557):871–876, Aug. 2021. Publisher: American Association for the Advancement of Science.

[3] M. L. Chaillet, S. Roet, R. C. Veltkamp, and F. Förster. pytom-match-pick: A tophat-transform constraint for automated classification in template matching. Journal of Structural Biology: X, 11:100125, 2025.

[4] J. Cheng, B. Li, L. Si, and X. Zhang. Determining structures in a native environment using single-particle cryoelectron microscopy images. The Innovation, 2(4):100166, Nov. 2021.

[5] J. Cheng, C. Wu, J. Li, Q. Yang, M. Zhao, and X. Zhang. Capturing eukaryotic ribosome dynamics in situ at high resolution. Nature Structural & Molecular Biology, 32(4):698–708, Apr. 2025. Publisher: Nature Publishing Group.

[6] S. Colvin, E. Jolibois, H. Ramezani, A. Garcia Badaracco, T. Dorsey, D. Montague, S. Matveenko, M. Trylesinski, S. Runkle, D. Hewitt, A. Hall, and V. Plot. Pydantic Validation, June 2025.

[7] J. L. Dickerson, E. Leahy, M. J. Peet, K. Naydenova, and C. J. Russo. Accurate magnification determination for cryoEM using gold. Ultramicroscopy, 256:113883, Feb. 2024.

[8] J. L. Dickerson and B. A. Lucas. To tilt or not to tilt? Strategies for in situ cryo-EM data collection. Current Opinion in Structural Biology, 93:103100, Aug. 2025.

[9] J. Elferich, L. Kong, X. Zottig, and N. Grigorieff. Ctffind5 provides improved insight into quality, tilt and thickness of tem samples. eLife, Nov. 2024.

[10] T. Grant and N. Grigorieff. Measuring the optimal exposure for single particle cryo-EM using a 2.6 Å reconstruction of rotavirus VP6. eLife, 4:e06980, 2015.

[11] M. Groll, L. Ditzel, J. Löwe, D. Stock, M. Bochtler, H. D. Bartunik, and R. Huber. Structure of 20S proteasome from yeast at 2.4Å resolution. Nature, 386(6624):463–471, Apr. 1997. Publisher: Nature Publishing Group.

[12] C. R. Harris, K. J. Millman, S. J. van der Walt, R. Gommers, P. Virtanen, D. Cournapeau, E. Wieser, J. Taylor, S. Berg, N. J. Smith, R. Kern, M. Picus, S. Hoyer, M. H. van Kerkwijk, M. Brett, A. Haldane, J. F. del Río, M. Wiebe, P. Peterson, P. Gérard-Marchant, K. Sheppard, T. Reddy, W. Weckesser, H. Abbasi, C. Gohlke, and T. E. Oliphant. Array programming with NumPy. Nature, 585(7825):357–362, Sept. 2020.

[13] R. Henderson. The potential and limitations of neutrons, electrons and x-rays for atomic resolution microscopy of unstained biological molecules. Quarterly Reviews of Biophysics, 28(2):171–193, 1995.

[14] J. Jumper, R. Evans, A. Pritzel, T. Green, M. Figurnov, O. Ronneberger, K. Tunyasuvunakool, R. Bates, A. Žídek, A. Potapenko, A. Bridgland, C. Meyer, S. A. A. Kohl, A. J. Ballard, A. Cowie, B. Romera-Paredes, S. Nikolov, R. Jain, J. Adler, T. Back, S. Petersen, D. Reiman, E. Clancy, M. Zielinski, M. Steinegger, M. Pacholska, T. Berghammer, S. Bodenstein, D. Silver, O. Vinyals, A. W. Senior, K. Kavukcuoglu, P. Kohli, and D. Hassabis. Highly accurate protein structure prediction with AlphaFold. Nature, 596(7873):583–589, July 2021. Publisher: Nature Publishing Group.

[15] A. A. Korostelev. The Structural Dynamics of Translation. Annual Reviews of Biochemistry, 91:245–267, June 2022. Publisher: Annual Reviews.

[16] B. A. Lucas. Visualizing everything, everywhere, all at once: Cryo-EM and the new field of structureomics. Current Opinion in Structural Biology, 81:102620, Aug. 2023.

[17] B. A. Lucas and N. Grigorieff. Quantification of gallium cryo-FIB milling damage in biological lamellae. Proceedings of the National Academy of Sciences, 120(23):e2301852120, June 2023. Publisher: Proceedings of the National Academy of Sciences.

[18] B. A. Lucas, B. A. Himes, and N. Grigorieff. Baited reconstruction with 2D template matching for high-resolution structure determination in vitro and in vivo without template bias, Oct. 2023. Pages: 2023.07.03.547552 Section: New Results.

[19] B. A. Lucas, B. A. Himes, L. Xue, T. Grant, J. Mahamid, and N. Grigorieff. Locating macromolecular assemblies in cells by 2D template matching with cisTEM. eLife, 10, June 2021. Publisher: eLife Sciences Publications Ltd.

[20] B. A. Lucas, K. Zhang, S. Loerch, and N. Grigorieff. In situ single particle classification reveals distinct 60S maturation intermediates in cells. eLife, 11, Aug. 2022. Publisher: eLife Sciences Publications, Ltd.

[21] V. J. Maurer, L. Grunwald, D. M. Kennes, and J. Kosinski. Constrained template matching using rejection sampling. bioRxiv, 2025.

[22] V. J. Maurer, M. Siggel, and J. Kosinski. Pytme (python template matching engine): A fast, flexible, and multi-purpose template matching library for cryogenic electron microscopy data. SoftwareX, 25:101636, 2024.

[23] T. pandas development team. pandas-dev/pandas: Pandas, Feb. 2020.

[24] A. Paszke, S. Gross, F. Massa, A. Lerer, J. Bradbury, G. Chanan, T. Killeen, Z. Lin, N. Gimelshein, L. Antiga, A. Desmaison, A. Köpf, E. Yang, Z. DeVito, M. Raison, A. Tejani, S. Chilamkurthy, B. Steiner, L. Fang, J. Bai, and S. Chintala. PyTorch: An Imperative Style, High-Performance Deep Learning Library, Dec. 2019. arXiv:1912.01703 [cs].

[25] E. F. Pettersen, T. D. Goddard, C. C. Huang, E. C. Meng, G. S. Couch, T. I. Croll, J. H. Morris, and T. E. Ferrin. UCSF ChimeraX: Structure visualization for researchers, educators, and developers. Protein Science : A Publication of the Protein Society, 30(1):70, Jan. 2021. Publisher: Wiley-Blackwell.

[26] J. P. Rickgauer, H. Choi, J. Lippincott-Schwartz, and W. Denk. Label-free single-instance protein detection in vitrified cells. bioRxiv, page 2020.04.22.053868, Jan. 2020.

[27] J. P. Rickgauer, H. Choi, A. S. Moore, W. Denk, and J. Lippincott-Schwartz. Structural dynamics of human ribosomes in situ reconstructed by exhaustive high-resolution template matching. Molecular Cell, 84(24):4912–4928.e7, 2024.

[28] J. P. Rickgauer, N. Grigorieff, and W. Denk. Single-protein detection in crowded molecular environments in cryo-EM images. eLife, 6:e25648, 2017.

[29] P. B. Rosenthal and R. Henderson. Optimal Determination of Particle Orientation, Absolute Hand, and Contrast Loss in Single-particle Electron Cryomicroscopy. Journal of Molecular Biology, 333(4):721–745, Oct. 2003. Publisher: Academic Press.

[30] S. H. Scheres. Relion: Implementation of a bayesian approach to cryo-em structure determination. Journal of Structural Biology, 180(3):519–530, 2012.

[31] F. J. Sigworth. Classical detection theory and the cryo-EM particle selection problem. Journal of Structural Biology, 145:111–122, 2004.

[32] E. Svidritskiy, A. F. Brilot, C. S. Koh, N. Grigorieff, and A. A. Korostelev. Structures of Yeast 80S Ribosome-tRNA Complexes in the Rotated and Nonrotated Conformations. Structure, 22(8):1210–1218, Aug. 2014. Publisher: Cell Press.

[33] P. Tesina, E. Heckel, J. Cheng, M. Fromont-Racine, R. Buschauer, L. Kater, B. Beatrix, O. Berninghausen, A. Jacquier, T. Becker, and R. Beckmann. Structure of the 80S ribosome–Xrn1 nuclease complex. Nature Structural and Molecular Biology, 26:275–280, 2019.

[34] K. Tunyasuvunakool, J. Adler, Z. Wu, T. Green, M. Zielinski, A. Žídek, A. Bridgland, A. Cowie, C. Meyer, A. Laydon, S. Velankar, G. J. Kleywegt, A. Bateman, R. Evans, A. Pritzel, M. Figurnov, O. Ronneberger, R. Bates, S. A. A. Kohl, A. Potapenko, A. J. Ballard, B. Romera-Paredes, S. Nikolov, R. Jain, E. Clancy, D. Reiman, S. Petersen, A. W. Senior, K. Kavukcuoglu, E. Birney, P. Kohli, J. Jumper, and D. Hassabis. Highly accurate protein structure prediction for the human proteome. Nature, 596(7873):590–596, Aug. 2021. Publisher: Nature.

[35] X. You, X. Zhang, J. Cheng, Y. Xiao, J. Ma, S. Sun, X. Zhang, H.-W. Wang, and S.-F. Sui. In situ structure of the red algal phycobilisome–PSII–PSI–LHC megacomplex. Nature, 616(7955):199–206, Apr. 2023. Publisher: Nature Publishing Group.

[36] K. Zhang, P. Cossio, A. V. Rangan, B. A. Lucas, and N. Grigorieff. A new statistical metric for robust target detection in cryo-EM using 2D template matching. IUCrJ, 12(2), Mar. 2025. Number: 2 Publisher: International Union of Crystallography.

[37] S. Q. Zheng, E. Palovcak, J.-P. Armache, K. A. Verba, Y. Cheng, and D. A. Agard. Motioncor2: anisotropic correction of beam-induced motion for improved cryo-electron microscopy. Nature Methods, 14(4):331–332, Apr 2017.

[38] W. Zheng, Y. Zhang, J. Wang, S. Wang, P. Chai, E. J. Bailey, W. Guo, S. C. Devarkar, S. Wu, J. Lin, K. Zhang, J. Liu, I. B. Lomakin, and Y. Xiong. Visualizing the translation landscape in human cells at high resolution. bioRxiv, 2024.

